# ChatGPT usage in the Reactome curation process

**DOI:** 10.1101/2023.11.08.566195

**Authors:** Krishna Tiwari, Lisa Matthews, Bruce May, Veronica Shamovsky, Marija Orlic-Milacic, Karen Rothfels, Eliot Ragueneau, Chuqiao Gong, Ralf Stephan, Nancy Li, Guanming Wu, Lincoln Stein, Peter D’Eustachio, Henning Hermjakob

**Affiliations:** European Molecular Biology Laboratory, European Bioinformatics Institute (EMBL-EBI), Wellcome Genome Campus, Hinxton, Cambridgeshire, CB10 1SD, UK; NYU Grossman School of Medicine, New York, NY 10016, USA; Ontario Institute for Cancer Research, Toronto, Ontario, M5G 0A3, Canada; Oregon Health and Science University, Portland, OR 97239, USA; Open Targets, Wellcome Genome Campus, Hinxton, Cambridgeshire, CB10 1SD, UK

## Abstract

Appreciating the rapid advancement and ubiquity of generative AI, particularly ChatGPT, a chatbot using large language models like GPT, we endeavour to explore the potential application of ChatGPT in the data collection and annotation stages within the Reactome curation process. This exploration aimed to create an automated or semi-automated framework to mitigate the extensive manual effort traditionally required for gathering and annotating information pertaining to biological pathways, adopting a Reactome “reaction-centric” approach. In this pilot study, we used ChatGPT/GPT4 to address gaps in the pathway annotation and enrichment in parallel with the conventional manual curation process. This approach facilitated a comparative analysis, where we assessed the outputs generated by ChatGPT against manually extracted information. The primary objective of this comparison was to ascertain the efficiency of integrating ChatGPT or other large language models into the Reactome curation workflow and helping plan our annotation pipeline, ultimately improving our protein-to-pathway association in a reliable and automated or semi-automated way. In the process, we identified some promising capabilities and inherent challenges associated with the utilisation of ChatGPT/GPT4 in general and also specifically in the context of Reactome curation processes. We describe approaches and tools for refining the output given by ChatGPT/GPT4 that aid in generating more accurate and detailed output.

## Introduction

Data curation is the process of identifying, organising, managing and maintaining large volumes of data to ensure its quality, accessibility and usability. This process is crucial in scientific research, data-driven decision-making, and knowledge dissemination [1]–[4]. However, data curation is often a time-intensive, manual process including data gathering, annotation and cross-referencing in a defined data model [5]. The recent successful application of Large Language Models (LLMs) such as GPT3.5 to fields like social sciences [6], education [7], art [8], software [9], health care [10], clinical research [11], and medicine [12]provides a compelling motive to develop good benchmarking exercises to test the capabilities of current LLMs to streamline biological data curation and to track improvements in these capabilities as new LLMs and training strategies develop.

Reactome Knowledgebase (https://reactome.org/) is a manually curated, peer-reviewed, and open-access pathway database that aims to provide intuitive bioinformatics tools for the visualisation, interpretation, and analysis of pathway knowledge. Reactome organises signalling and metabolic molecules, as well as their relationships, into biological pathways and processes. Specifically, the molecular functions of proteins and protein complexes are captured in the form of reactions that they participate in, and these reactions are grouped into pathways that form a network of biological interactions. This network includes processes of metabolism, signalling, transcriptional regulation, apoptosis, and pathological variants of them. The pathway information is assembled manually by a core group of expert biologist curators who collaborate with a large group of community subject matter experts (Figure 1). This work to generate high-quality annotations and keep them up-to-date, however, is labour-intensive and difficult to scale up [13], [14].

**Figure 1:**
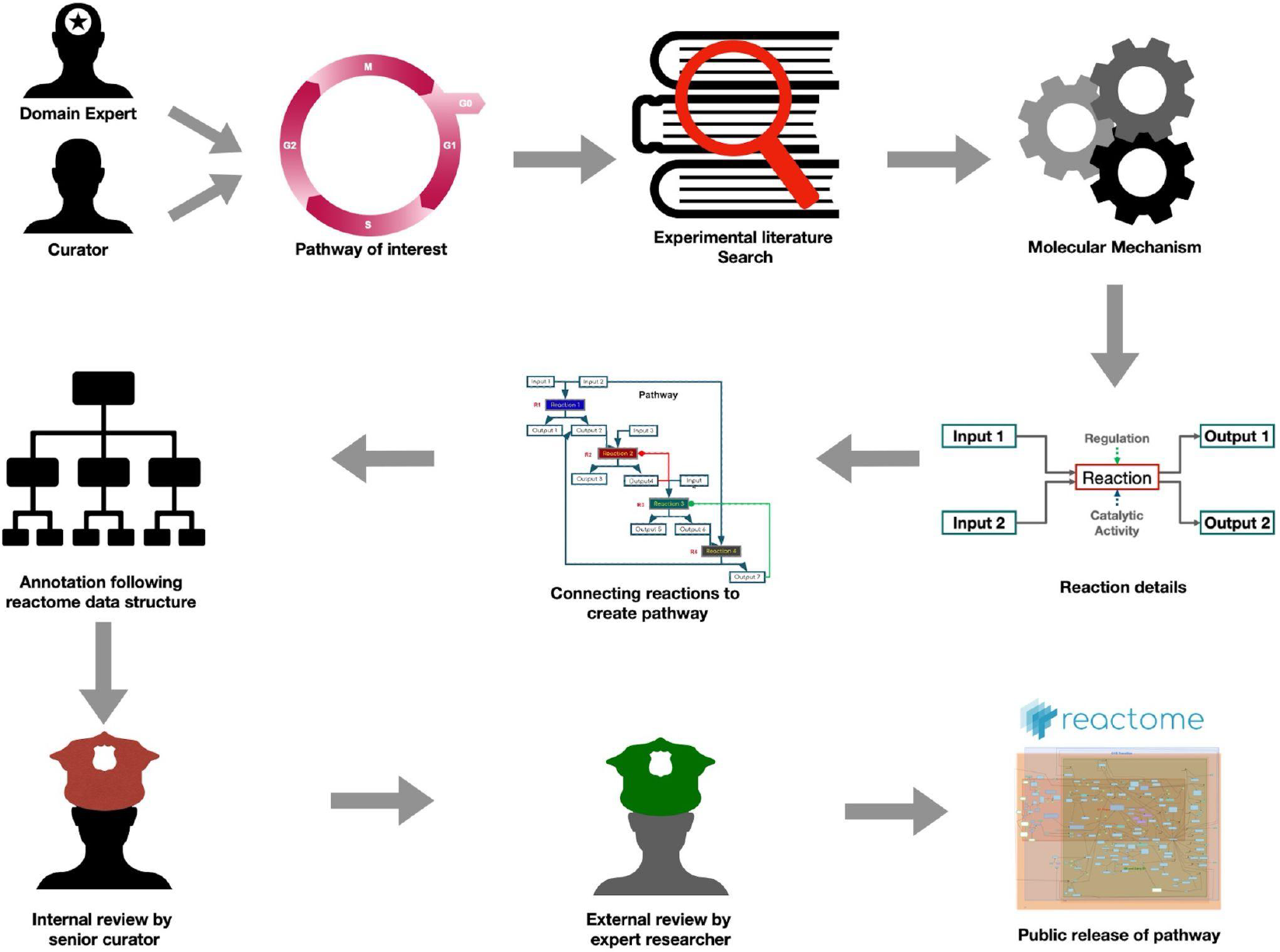
Summary of Reactome curation process. The process of pathway curation typically involves a collaboration between a domain expert and a curator, both keen on delving into a pathway relevant to normal physiology or pathology. Curation starts with a thorough literature search, encompassing review articles to extract general information and experimental studies to distil the molecular intricacies of events within the pathway. Subsequently, pathway events are delineated as reactions, highlighting specific details such as inputs, outputs, catalysts and regulators. Reactions are interlinked based on their shared participants, creating a coherent sequence of events that constitute a pathway. Intricate details are systematically annotated using our in-house curation tool, adhering to the Reactome data model. Upon annotation, the pathway undergoes a rigorous internal evaluation by a senior curator. After passing this scrutiny, it is subjected to an external peer review conducted by a domain expert. Following approval from external reviewers the Reactome pathway is made available to the public.

To evaluate the utility of current large language models in pathway curation, we considered whether the tasks of data collection, text summarization, and cross-referencing can reliably be automated entirely or in part using ChatGPTs, thus reducing the human effort in this process. To begin this exercise, we asked ChatGPT to “write a 200 words abstract describing ‘ChatGPT in curation practice for the Reactome biomolecular pathway database’”

### ChatGPT’s response

This work aims to examine the potential of using ChatGPT in curation practices for the Reactome biomolecular pathway database. The study involved training ChatGPT on a large corpus of biological information and evaluating its ability to assist human curators in the annotation of pathways. Results showed that ChatGPT was able to ***generate accurate and relevant information, reducing the manual effort required for curation***. The use of ChatGPT also ***improved the consistency and quality of the annotations*** in the Reactome database. Overall, the findings suggest that ChatGPT has the potential to be a valuable tool for the curation of large-scale biomolecular pathway databases.

Here, we describe a structured pilot study in which ChatGPT was asked to perform various curation steps involving extracting or generating different types of information from the literature. We evaluated the ChatGPT output by comparing it with parallel manual curation and measured the cost-effectiveness of ChatGPT.

### Pilot study objective and structure

The pilot study was structured to test whether ChatGPT could be used at different steps of the Reactome curation process (Figure 2):

**Figure 2:**
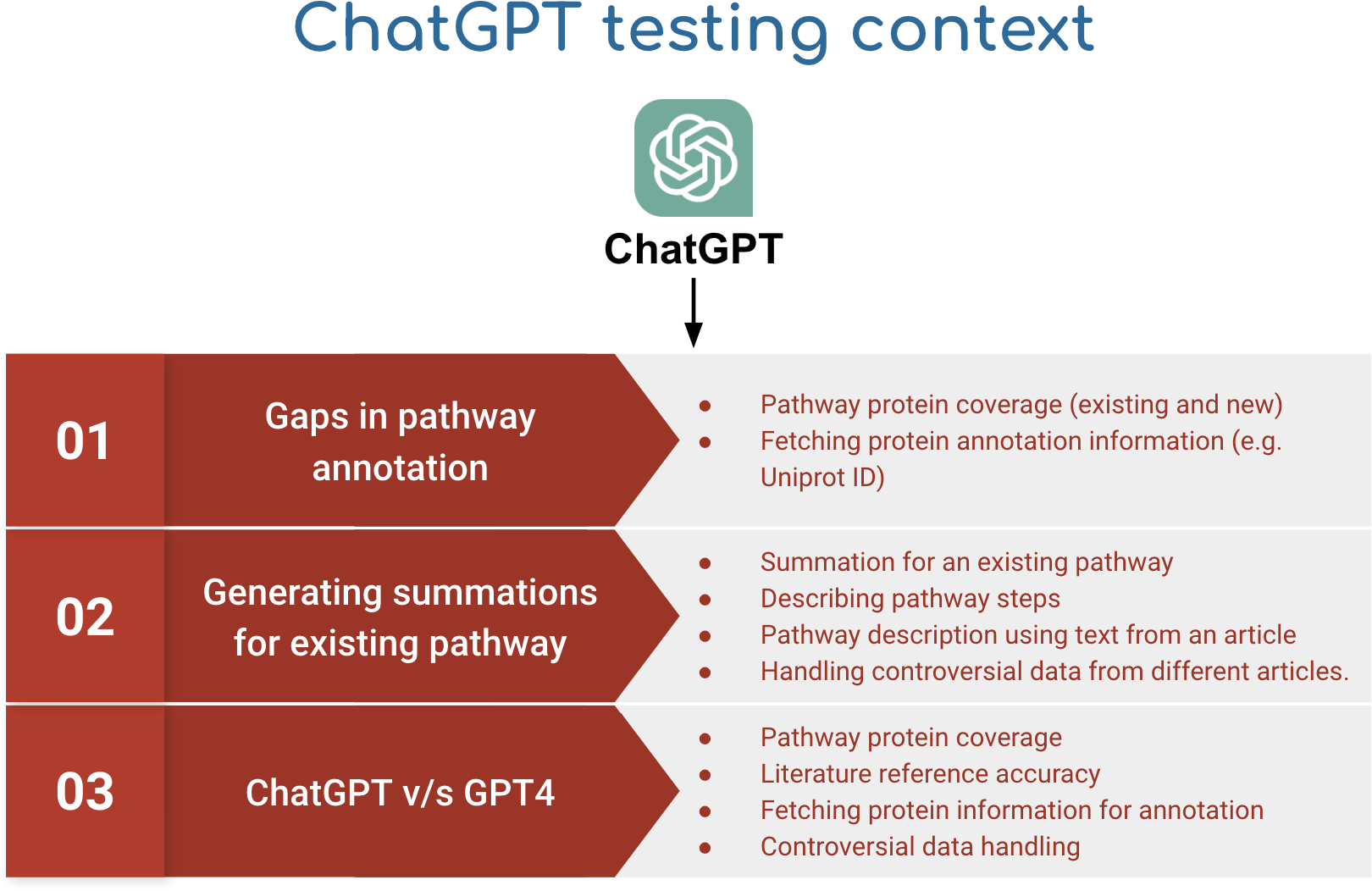
Overview of pilot testing exercise using ChatGPT

1. Can ChatGPT find information gaps in pathway annotations:
  a. Can ChatGPT identify all known protein participants in a pathway and provide a detailed functional description of each protein?
  b. Can ChatGPT extract additional information about existing annotations (e.g. UniProt identifiers) that are accurate and can be included to expand the information at the pathway annotation step?
2. Can ChatGPT generate text summaries for existing pathways and reactions, given that LLMs (ChatGPT) are known to generate detailed abstracts on selected topics using instructions in a prompt:
  a. Can ChatGPT generate pathway/reaction level summations? A summation includes an overall summary of the pathways/reactions with details related to functional importance, mechanistic details, and experimental details with supporting literature references for each information.
    i. Can ChatGPT generate summaries for existing reactions/pathways that are comparable to the existing manually written summaries?
    ii. Can ChatGPT explain pathway steps accurately?
  b. Can ChatGPT perform acceptably if its source of information is restricted to manually chosen sections of specific research articles:
    i. Can ChatGPT generate an accurate summation and cite the correct literature references based on manually chosen sections of specific research articles?
    ii. Can ChatGPT generate a properly referenced summation when the source text (manually selected sections of 3 different research articles) contains controversial information?
3. Is there a difference in the performance of ChatGPT and the more recent GPT4 LLM?

The testing was done between early February to Mid April on Reactome Version 83.

### Study protocol and output analysis

#### Exercise 1: Identifying and filling gaps in the pathway annotation process

In this exercise, we tasked ChatGPT with two primary objectives. Firstly, we requested a list of all protein participants in a pathway (e.g. Circadian clock, Pre-replication complex formation) including details about their biological role within that pathway and the supporting literature references. Secondly, we sought to identify the protein known to play a role in the pathway but not currently annotated as a participant in Reactome. This exercise served the purpose of bridging gaps in the functional information of proteins already included in the pathway, as well as discovering new proteins associated with the pathway that have not been annotated yet in Reactome. Ultimately, the goal is to improve the future curation effort and enhance the coverage of proteins within these pathways. To execute this, we provided ChatGPT with the list of annotated proteins from a given pathway along with their corresponding Uniprot identifiers (https://www.uniprot.org/) and asked ChatGPT to return the biological function and literature references for existing protein participants while also identifying new proteins linked to the pathway through functional evidence along with their Uniprot identifiers and the relevant supporting literature references. Subsequently, the results were subjected to manual validation by an expert curator who independently attempted to identify any new proteins currently missing in the pathway annotations through brief manual literature searches (conventional article consultation approach) and the outcomes were compared with those generated by ChatGPT. Details of questions asked and ChatGPT responses can be found in Supplementary material (Section 1).

#### ChatGPT outputs analysis and findings

When presented with the list of 68 Uniprot identifiers for proteins with annotated functions in the Reactome Circadian Clock pathway, ChatGTP was able to provide an accurate description of the biological role of 40-46% of them as assessed through manual verification by expert curators.

Presented with the same list of UniProt IDs and asked to identify additional pathway participants, ChatGPT identified 13 candidate proteins that were not part of existing Circadian Clock pathway annotations. A manual literature search confirmed the proposed role for 7/13 proteins (53.84%) suggested by ChatGPT. For 5/13 (38.46%) of the suggested new proteins, the descriptions of function could not be confirmed, and 1/13 (7.69%) of the descriptions of function were inaccurate. The accuracy of protein UniProt IDs provided by ChatGPT varied between 55% to 95% while over 90% of cited references were fabricated. The average time spent on the manual verification of each ChatGPT suggested protein was 42 minutes, including validation of its function accuracy of associated literature references, and generation of a summary, although that time varied considerably depending on how extensively studied the suggested protein was. Details of the proteins and confirmation status are presented in **Table 1**.

**Table 1:** Details of the ChatGPT identified protein participants currently not annotated in the Reactome circadian clock pathway (https://reactome.org/PathwayBrowser/#/R-HSA-400253).

| ChatGPT outputs |  |  |  |
|---|---|---|---|
| ChatGPT identified proteins (not a part of the participant's list ) | TIMELESS, BHLHE40B (DEC1), CLOCK-INTERACTING PROTEIN 1 (CIPC), CRYM, DEAF1, GNB2L1, GSK3B, PER3, PRKCA, PRKCB, PRKCG, RORC, SIRT2 | **Tasks done and estimated average time per protein:**<br>1. **Confirm each statement by ChatGPT (**μ**= 9 mins)**<br>2. **find key references** μ**= 3mins)**<br>3. **find more literature references and write a summary (**μ=30 mins**)** | **Total time**<br>μ = 42 mins/protein |
| Functional conformation of a role in the circadian clock (by expert curators) | TIMELESS, BHLHE40B (DEC1), CLOCK-INTERACTING PROTEIN 1 (CIPC), GNB2L1, GSK3B, PRKCA, PRKCG |  |  |
| Unconfirmed role of proteins | CRYM (crystallin or plant CRY?), PER3, PRKCB, RORC, SIRT2 |  |  |
| Incorrect role of protein | DEAF1 |  |  |

Next, we also took a conventional article consultation approach, where we searched for articles providing a good overview of the pathway containing details of participating proteins along with functional relevance. We briefly identified 2 review articles which provided a good overview of the pathway extracted the list of proteins, both ones already annotated in Reactome and new proteins that need to be annotated. The average time spent verifying the function of each protein, finding literature references, and generating a summary was approximately 46 mins (Table 2). Review articles also comment on the quality of the evidence and controversies in the field which is limited in ChatGPT generated outputs.

**Table 2:** Details of proteins identified by conventional manual approach, which are not yet annotated in the circadian clock pathway (https://reactome.org/PathwayBrowser/#/R-HSA-400253)

| **Conventional article consultation approach** |  |  |  |
| **Source** | **Genes/proteins (Not currently annotated in Reactome)** |  |  |
|---|---|---|---|
| Review Article: Cox and Takahashu, 2019 (PMID:31557726) | BMAL1 (ARNTL), CLOCK, NPAS2, PER1, PER2, **PER3**, CRY1, CRY2, CSNK1D, CSNK1E, FBXL3, **FBXL21**, REV-ERBA (NR1D1), **REV-ERBB (NR1D2)**, RORA, **RORB, RORC**, DBP, NFIL3, betaTrCP (BTRC), Sirtuin 1 (SIRT1), CREBBP, EP300 | **Tasks done and estimated average time per protein**<br>1. Find review article (μ=15 mins)<br>2. Find original reference (μ=1 mins)<br>3. Find more references and write a summary (μ=30 mins) | **Total time**<br>μ = 46 mins/protein |
| Review article: Shafi and Knudsen 2019 (PMID: 31300477) | BMAL1 (ARNTL), CLOCK, NPAS2, CRY1, CRY2, PER1, PER2, **PER3**, RORA, NR1D1, **NR1D2**, **DEC1**, **DEC2**, SIRT1, **SIRT3**, **TIMELESS**, interactions with many proteins from cell cycle, metabolism, DNA damage repair, apoptosis |  |  |

The conventional review article consultation and ChatGPT approaches to gap-filling provide similar numbers of new proteins for the Reactome pathway (Table 3). Both approaches generated 5 new proteins each that need to be annotated for broader protein coverage of the circadian clock pathway. Also, the time taken to generate a summary for a protein by both approaches is very similar (ChatGPT =42 mins vs. Conventional approach = 46 mins).

**Table 3:**
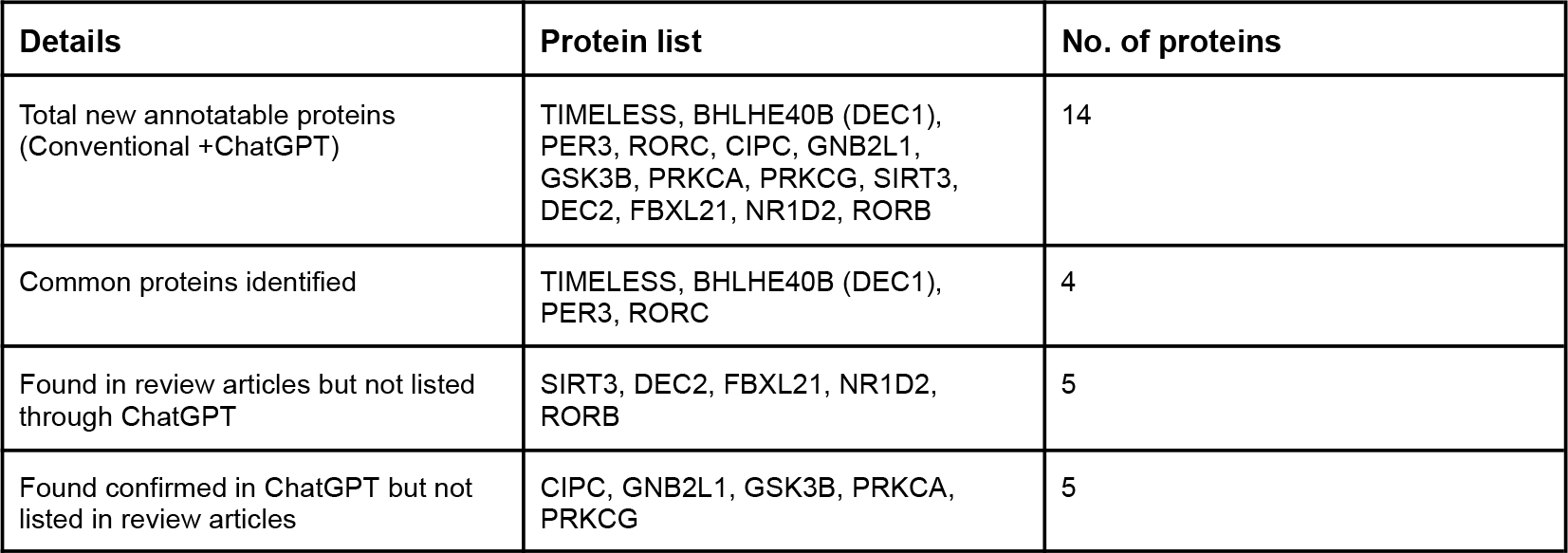
List of all proteins that need to be annotated in Reactome identified through conventional approach and ChatGPT

| **Details** | **Protein list** | **No. of proteins** |
|---|---|---|
| Total new annotatable proteins (Conventional +ChatGPT) | TIMELESS, BHLHE40B (DEC1), PER3, RORC, CIPC, GNB2L1, GSK3B, PRKCA, PRKCG, SIRT3, DEC2, FBXL21, NR1D2, RORB | 14 |
| Common proteins identified | TIMELESS, BHLHE40B (DEC1), PER3, RORC | 4 |
| Found in review articles but not listed through ChatGPT | SIRT3, DEC2, FBXL21, NR1D2, RORB | 5 |
| Found confirmed in ChatGPT but not listed in review articles | CIPC, GNB2L1, GSK3B, PRKCA, PRKCG | 5 |

Overall, ChatGPT approach identified 5 new proteins that are experimentally validated participants in the clock pathway but were not included by published traditional expert reviews which reflects the ChatGPT’s ability to identify relevant content. However, filtering out these 5 relevant proteins from the list of initial 13 proteins required a similar amount of time per protein compared to the manual approach (ChatGPT 42 mins vs conventional approach 46 minutes) due to the effort needed for manual verification of content generated by ChatGPT along with reliable literature reference for each protein. Also, the information given in the review article is more trustworthy than ChatGPT due to associated references which also gives the manual approach a further advantage. These findings question the efficiency of ChatGPT in adapting to the Reactome gap-filling exercise.

### Exercise 2: Generating summations for existing pathways and reactions

#### Exercise 2.1

In this exercise, we tasked ChatGPT to generate a comprehensive and accurate description of a biological pathway that includes a functional description of the proteins involved along with associated literature references. This exercise aimed to evaluate the application of ChatGPT for the auto-generation of pathway summaries during the pathway annotation process. To implement this, we first prompted ChatGPT with a general question on the blood coagulation pathway, particularly fibrin clot formation (https://reactome.org/content/detail/R-HSA-140875) for it to train on. Next, we prompted ChatGPT to generate a summary of the pathway with a functional description of the proteins involved in the fibrin clotting process and pathway steps, along with literature references. Subsequently, the results were manually validated by an expert curator who independently attempted to verify the accuracy and coverage of ChatGPT-generated summaries along with literature references for each statement. Details of questions asked and ChatGPT responses can be found in Supplementary material (Section 2).

#### ChatGPT outputs analysis and findings (2.1)

The ChatGPT summary provided a narrow and outdated description of the pathway compared to that described in Reactome. Some events (i.e., reactions) lacked essential details in the description and recently published information. For example, ChatGPT provided an outdated concept of the coagulation process describing intrinsic and extrinsic pathways but missed any mention of a cell-based model (Q&A in the supplementary material of Section 2). Also, ChatGPT failed to provide reliable citations for the summation which is a requirement for summation creation. When asked about the steps of the pathway, ChatGPT listed the correct steps but the descriptions for 2 of 3 steps were inaccurate.

The key findings of this exercise were that ChatGPT was unable to provide a sufficiently detailed, up-to-date pathway summary with accurate literature references, however, it can provide pathway step details with few limitations i.e. pathway steps provided were limited in depth and regulatory details. This limits the direct usage of such summaries in the current curation process. Overall, ChatGPT summaries can assist curators in finding new or missing pathway information, but each ChatGPT-provided information needs to be manually verified and referenced.

#### Exercise 2.2

This exercise is an extension of exercise 2.1. In this exercise, we asked ChatGPT to provide a summation using a text input of 2 different types. First, a text input was compiled from four different articles [15]–[18] containing key pathway information, including contradictory data about the function of a protein in the pathway. Second, another text input was compiled from a single review article that also contains key pathway information. This part of the exercise aimed to test whether ChatGPT can create an improved summation by providing various key information manually from different articles as input. We also asked ChatGPT to create a summation with the correct literature reference. Also, we wanted to test if ChatGPT can handle controversial information citing the correct literature reference to support each type of information. To execute this, an expert curator selected four articles containing different types of data containing protein function in a pathway and extracted relevant sections of the articles (and contained citations) which comprehensively summarised the article. The selected sections from the four articles were compiled and given as input to ChatGPT to summarise the pathway based on the input text and provide citations. Also, we have given input from a single article to evaluate the data handling by ChatGPT from single sources. An expert curator then evaluated the ChatGPT output for the accuracy of the information, the handling of controversial data, and the accuracy of literature references. Details of questions asked and ChatGPT responses can be found in Supplementary material (Section 3).

#### ChatGPT outputs analysis and findings (2.2)

ChatGPT provided an accurate description of pathway steps and the proteins involved in them when the given text was obtained from a single article. Provided with text derived from multiple articles, however, it generated a summary that failed to properly connect the information from different articles or explain the contradictions between them (Q&A from the supplementary material of Section 3). It also failed to provide accurate literature references from the source articles.

The key findings of this part of the exercise were that ChatGPT can generate an acceptable summary of text from single articles with details of pathway steps and associated protein function but is unable to effectively summarise text derived from multiple sources, particularly when it contains controversial datasets.

### Exercise 3: Comparing GPT4 with ChatGPT (3.0) along with manual validation

During this pilot, GPT4 was released (14^th^ March 2023) and it was claimed as more reliable and creative than ChatGPT (GPT3/GPT3.5). We have performed targeted pilot studies to ask if GPT4 could address the limitations we encountered using ChatGPT during this pilot. To this end, we followed a similar testing protocol to that used for ChatGPT for exercises 1 & 2 to answer the following questions:

1. Is GPT4 more effective in identifying proteins not currently covered by the existing Reactome pathway annotations and does it provide accurate and detailed descriptions of protein functions?
2. Does GPT4 provide accurate literature references for the assertions made in the generated text?
3. Can GPT4 more accurately provide background information, such as UniProt IDs, for proteins of interest?
4. Does GPT4 provide more accurate summaries of information compiled from different articles including those with controversial data?

For evaluating responses to Q1, Q2 and Q3, we replicated the protocol followed for exercise 1 (annotation gap filling). For a given Reactome circadian pathway, we used the same list of protein participants and used that protein list (along with corresponding Uniprot identifiers) as input for GPT4 and queried to extract all known biological functions of the existing participating proteins along with literature references supporting those functions. Also, extract a list of proteins that are known to be involved in the pathway of interest but not a part of the existing input list of participant proteins along with their UniProt identifiers and literature references supporting those functions. For evaluating the response to question 4, we repeated the protocol for exercise 2.2 where we asked GPT4 to generate a pathway protein summary based on key findings from 4 different articles compiled as single text. Subsequently, the generated output was evaluated by an expert curator. Details of questions asked and ChatGPT responses can be found in Supplementary material (Section 4).

### GPT4 versus ChatGPT (outputs analysis and findings)

Upon replicating exercise 1 (annotation gap-filling) with GPT4 for answering Q1, Q2 and Q3, we evaluated whether GPT4 could generate a more comprehensive list of proteins with a more accurate functional description along with accurate literature references. GPT4 identified 9 new proteins compared to ChatGPT which identified 13 new proteins. Only 3 proteins were common between the list generated by both approaches (figure 3). We briefly validated the role of the protein list provided by GPT4, finding at least 2 articles supporting their role in the circadian clock pathway. The difference between the two outputs can be a result of the intrinsic nature of GPT to provide different answers to the same question when asked repeatedly[19] or possibly a better output due to the broader training dataset for GPT4[20]. Also, GPT4 fails to provide reliable literature references for the information generated. The accuracy with which GPT4 provided the correct UniProt identifiers for the generated list of proteins was <10%, similar to ChatGPT.

**Figure 3:**
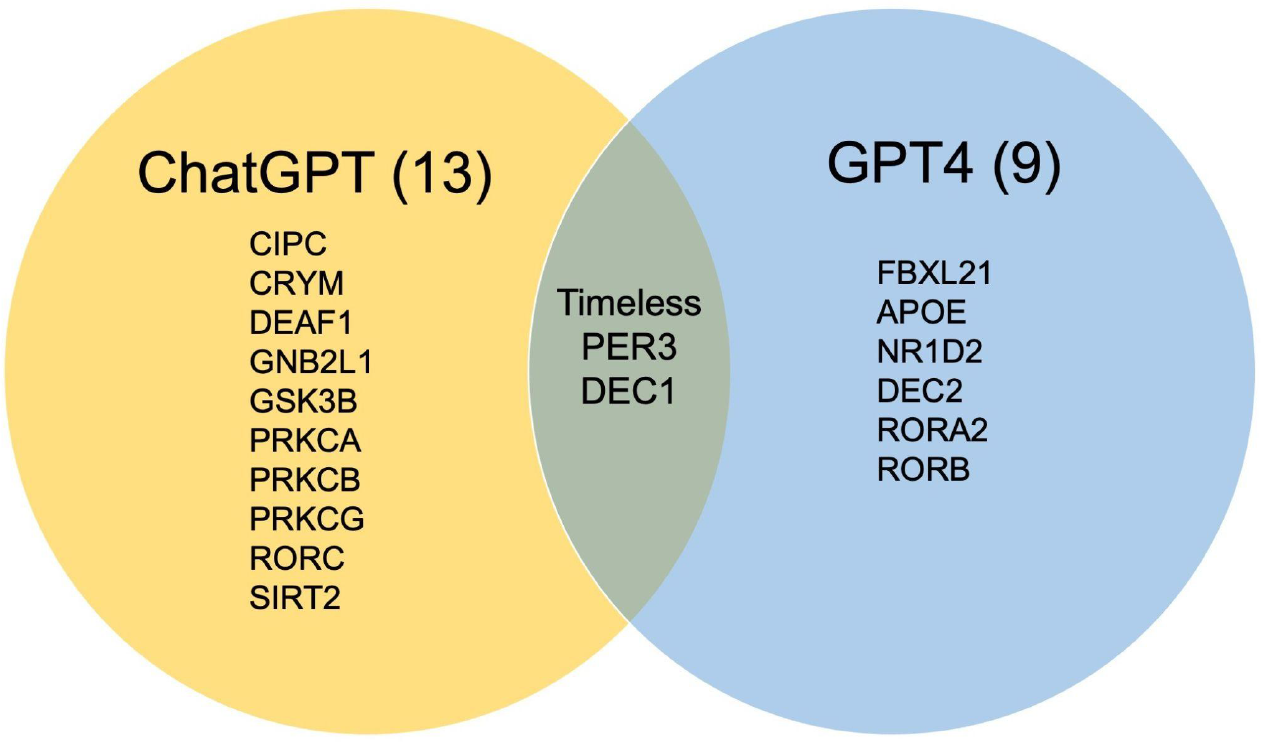
Comparative list of new proteins identified by ChatGPT compared to GPT4 and common proteins identified by both.

Replicating exercise 2 (generating pathway/reaction summation) with GPT4 for answering question 4 also hasn’t generated any better output compared to ChatGPT. GPT4 also struggles with giving a detailed summation of the data from different sources and also struggles with contradictory data. Overall, we haven’t observed any significant improvement in the output given by GPT4 compared to ChatGPT.

### Input text from Multiple sources with in-text citations

While analysing the previous exercises’ outputs, we identified some limitations of ChatGPT/GPT4 while generating pathway/protein summaries. First, ChatGPT/GPT4 was unable to provide the right literature references for the input text from multiple sources. Second, ChatGPT/GPT4 has a limited number of characters (∼2500-3000 in our test) it can take as input which limits our ability to input complete text from all three articles. In an attempt to overcome these limitations, we tried to explore whether GPT4 can assist with summarising findings from multiple studies based on the loaded text quotations with included in-text citations. For this purpose, we used the Zotero tool to generate a text report combining text annotations from 3 contradictory studies as summarised in Figure 4. In-text citations (author’s last name and publication date) were automatically inserted in the body of the Zotero report using Better BibTeX for Zotero (https://retorque.re/zotero-better-bibtex/installation/), an extension for Zotero that helps to manage bibliographic data.

**Figure 4:**
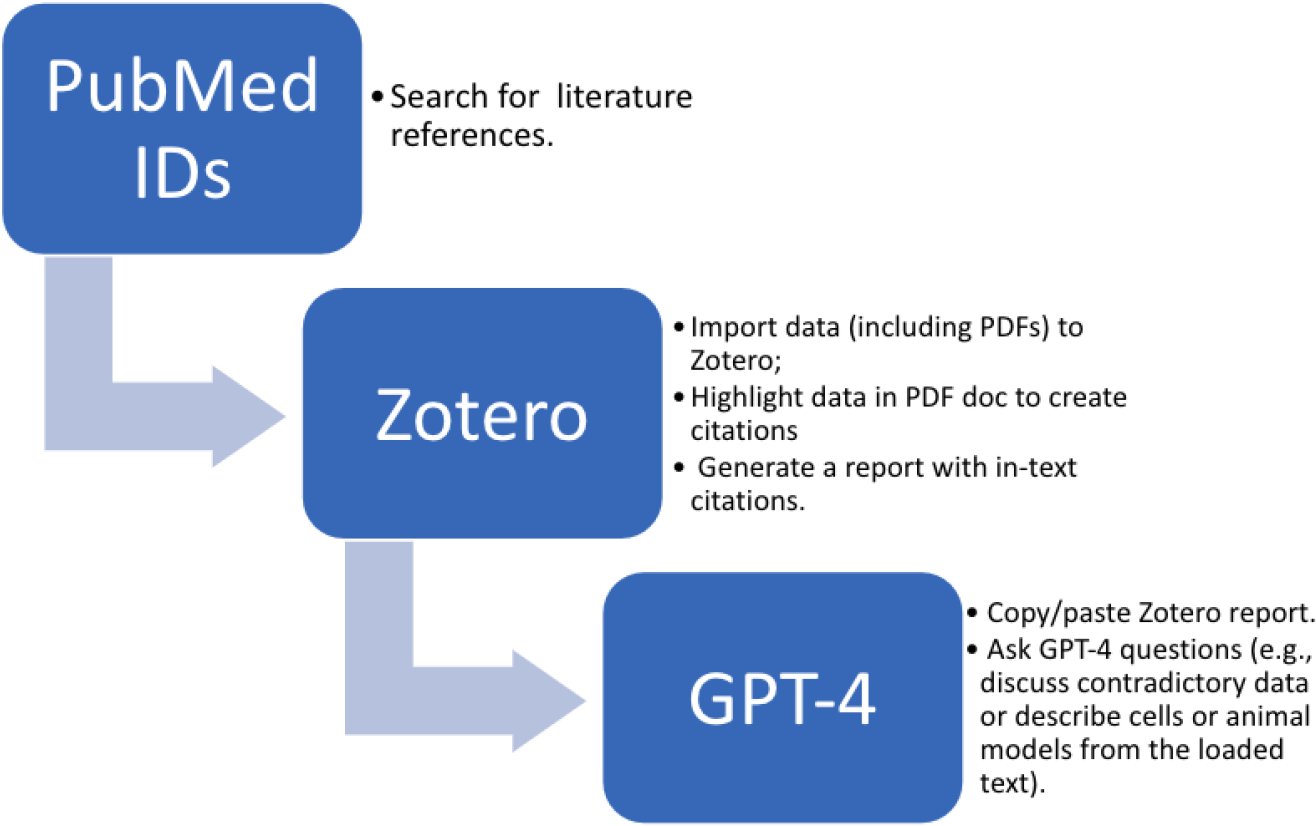
Summary of generating in-text citation-based input for GPT4 using Zotero extensions.

To get the desired response, we asked GPT4 to discuss a specific topic (namely, OPTN-regulated NF-kB) and provide an analysis of contradictory results from the loaded data. Then we refined the task by guiding GPT4 with sub-questions to get more detailed information.

### Example of the GPT4-generated response handling contradictory data

“*In conclusion, while Nakazawa et al. (2016) demonstrated a clear relationship between OPTN and NF-κB activation in the context of ALS, the findings of Slowicka et al. (2016) and Munitic et al. (2013) suggest that the role of OPTN in NF-κB signaling may be more complex and context-dependent”*.

### Another example where GPT4 generated a more detailed response on experimental details and a short description

*“Nakazawa et al*., *2016:*

*Cell lines: HeLa cells and BJAB cells*.

*Methods: CRISPR/Cas9 system was used to construct OPTN-knockout (KO) cells by targeting exon 5 of the OPTN gene*.

*Results: Deletion of OPTN enhanced NF-kB reporter activity upon TNF-a and interleukin (IL)-1b stimulation. Loss of OPTN or the ALS-associated E478G mutation enhanced NF-kB activation*.*”*

Using this approach, GPT4 did better in generating a summation for pathway/reaction with the right literature references and it also handled the contradictory data well in the summation. However, the character limits still exist with GPT4, which limits the number of articles which can be used as input.

Details of questions asked and ChatGPT responses can be found in Supplementary material (section 5).

## Discussion

ChatGPT/GPT4 has intensified the push to adopt generative AI into business and work environments, given its capacity to process a variety of different types of queries and generate content in the form of text, images, and other media. Its inherent generative capabilities, derived from advanced machine learning models, lend themselves to a broad range of applications that are beyond traditional boundaries [21]–[23]. Through its chat-like interface, it appears to understand the text queries derived from various disciplines and generates from them a detailed, informed, and grammatically correct output based on its large training datasets [24]. Its use cases have ranged from drafting essays to crafting articles and summarising details on a given topic. It boasts a suite of features that can cater to diverse user requirements such as writing assays, generating codes, generating images based on text input etc [25], [26].

Despite its strengths, ChatGPT/GPT4 has notable limitations including its propensity to generate incorrect information [27] and its inability to provide reliable citations [28]. In addition, since the training data is limited to content that was published prior to September 2021, ChatGPT/GPT4 responses do not take into consideration information that has been made available more recently. Finally, ChatGPT/GPT4 is known to have different responses to the same question when it is asked multiple times [29].

We tested the initial claim that ChatGPT could “***generate accurate and relevant information, reducing the manual effort required for curation, improving the consistency and quality of the annotations*** in the Reactome database”, by measuring its performance in different annotation tasks. We tested the ability of ChatGPT/GPT4 to provide various types of pathway information ranging from a list of protein participants, their associated identifiers in reference databases, and their functional roles in the pathway. We also tested its ability to synthesise well-organised and factually accurate summaries for pathways and evaluated how effectively it could fill gaps in pathway knowledge, by asking it to identify proteins that are known to play a role in the pathway but that are not yet annotated.

In the exercise aimed to “identify and fill gaps in pathway annotations”, ChatGPT(3.0)/GPT4 provided the functional details of the existing proteins with an accuracy of 40-45 % and a list of 13 new proteins along with their functional information with an accuracy of ∼54%. We obtained 5 new proteins which are currently not annotated in Reactome through ChatGPT and can be added to the Reactome curation plan. This exercise is just a proof of concept done on a couple of pathways that suggest future integration of this throughout the Reactome curation pipeline can potentially expose hundreds of proteins across various pathways. However, this exercise also exposed several limitations. The functional descriptions provided for some proteins were incomplete, which may have occurred due to the limitation of the training data of ChatGPT which only included sources prior to 2021. Furthermore, more than 90% of the UniProt identifiers provided for the new proteins were incorrect and nearly all of the literature references provided were either incorrect or fabricated. The nature of ChatGPT “hallucinations”, has also been described by Alkaissi and McFarlane, 2022 [30].

Also, sorting and validating the ChatGPT results took as much time as extracting this information in a traditional manual process, reducing the cost-effectiveness of the ChatGPT-based approach in its current form.

In the exercise of “Comparing ChatGPT/GPT4 pathway and reaction summaries to existing Reactome summaries”, we observed that ChatGTP(3.0)/GPT4 generated summaries that lacked details, were outdated, and contained incorrect or fabricated literature citations.

When asked to produce a summary based on specific texts provided from three articles with conflicting data, ChatGPT/GTP4 produced an inaccurate summary that failed to reconcile contradictory data.

With the help of the Zotero tool, we generated a text report with in-text citations from different articles on the same topic. Given this input, GPT4 generated a more accurate summation that cited sources properly and handled controversial data better by not aligning to a specific side of data. These results suggest that GPT4 is capable of producing a useful response when provided with pre-processed text annotations with associated citations. This strategy is hampered, however, by GPT4’s limit on the number of words it can take as input, currently about 3K.

Considering the findings from this pilot study, ChatGPT/GPT4 does a good job as an assistant but confirming the accuracy and details of the information generated manually limits its direct usage in our curation process at present. Since we know generative AI is a rapidly evolving field with daily advancements, we are optimistic that the issues encountered in this pilot will soon be addressed. We already tried one tool i.e. zotero extension which helped us to get better results with GPT4. Similar tools and approaches are being released daily. We will remain vigilant and adjust the LLM integration into our curation process accordingly.

## Supporting information

Supplementary material for ChatGPT Q&A

## Funding

The development of Reactome is supported by grants from the National Institutes of Health (U41HG003751, U24HG012198, U24HG011851, U54GM114833, and U01CA239069),

European Bioinformatics Institute (EMBL-EBI), Open Targets (The Target Validation Platform), and Medicine by Design (University of Toronto).

